# Assessing the performance of TRX and DUF148 antigens for detection of prepatent Guinea worm (*Dracunculus medinensis*) infection in dogs

**DOI:** 10.1101/2024.09.12.612594

**Authors:** Hassan Hakimi, Pabasara Weerarathne, Meriam N. Saleh, Raquel R. Rech, Richard Ngandolo, Philip Ouakou Tchindebet, Sidouin K. Metinou, Lucienne Tritten, Guilherme G. Verocai

## Abstract

Guinea worm (GW, *Dracunculus medinensis*) is a nematode that causes a painful and debilitating neglected tropical disease in humans. The GW Eradication Program has decreased human infections by >99% over the last 40 years. However, GW emergence in animal hosts, particularly dogs, has hampered eradication efforts. Currently, there is no method for diagnosing GW infection in animals during the prepatent period, before the adult female worms emerge. Previous works have identified two GW proteins, TRX and DUF148, as immunoreactive antigens with GW-positive human and dog sera. This study developed and validated indirect enzyme-linked immunosorbent assays (ELISA) using each antigen alone or in a combination of both antigens. Using serum samples from experimentally exposed dogs, TRX and DUF148 showed reactivity at 9- and 11-weeks post-exposure, respectively. In an experimentally infected ferret, TRX and DUF148 showed reactivity at 13- and 15-weeks post-exposure, respectively. These antigens were further validated using sera of dogs from endemic villages in Chad (n=47) and shelter dogs from the non-endemic United States (n=492). DUF148 showed better reactivity and sensitivity of 76.6.% in detecting GW infection in prepatent sera compared to TRX. However, DUF148 cross-reacted with one serum sample from *Brugia pahangi* experimental infection and several shelter dog sera. The anti-DUF148 titer was significantly higher in the shelter dogs positive for gastrointestinal nematodes than in negative dogs. To mitigate this cross-reaction, we produced 3 peptides of DUF148. Peptide 3 from the C-terminal was more reactive with prepatent sera and had a sensitivity of 83%; however, the specificity was not superior to DUF148 whole antigen. The antibody response to DUF148 in Chad dogs with the history of GW emergence waned overtime but was detectable until two years post-GW-emergence. Our findings could facilitate the development of diagnostic methods for early detection of GW infection in dogs in endemic countries.

**Authors summary:** Dracunculiasis or Guinea worm (GW) disease is a neglected tropical disease caused by the nematode *Dracunculus medinensis* and is targeted for eradication by the World Health Organization. The main challenge for the eradication program is the emergence of animal infections, especially dogs. Diagnostic tests are needed to find infected dogs during the prepatent period to better control infection and prevent the spread of GW. Previous studies have found two immunoreactive GW proteins, TRX and DUF148. In this study, we validated these antigens to detect infection in before GW emergence. Using sera of dogs from endemic areas of Chad, we found that DUF148 was more reactive and had promising sensitivity to detect the prepatent infection in an indirect ELISA assay. However, DUF148 also showed cross-reaction with some sera of dogs from the Unites States, a non-endemic area for GW. To mitigate this cross-reactivity, we performed ELISA using shorter peptides. We found peptide 3 that covers the C-terminal of the protein is the immunogenic part of DUF148. However, peptide 3 ELISA did not outperform whole antigen ELISA. This study confirms the applicability of DUF148 ELISA in detecting prepatent infections in dogs and could assist the GW Eradication Program.

## Introduction

Guinea worm (GW, *Dracunculus medinensis*) is a parasitic nematode that causes dracunculiasis or GW disease, a debilitating and neglected tropical disease in humans. Humans acquire the parasite by ingesting infected cyclopoid copepods in drinking water. The copepod serves as the intermediate host and contains the infective third-stage GW larvae (L3) [1, 2]. The global GW Eradication Program (GWEP), which was started in 1986 and led by the Carter Center, has reduced annual cases from an estimated 3.5 million human cases in Africa and Asia [3] to 14 cases [4] in sub-Saharan Africa in 2023. GW is the first human parasitic disease set for eradication. The eradication effort is based on primary health care, health education, and promoting the utilization of filters for drinking water to prevent ingestion of copepods [1, 2].

Despite being successful in reducing human cases, the emergence of GW in animals, especially domestic dogs, poses an extra challenge to GWEP and may delay eradication. Although animal infections by GW were recorded previously both naturally and experimentally, these infections have disappeared following the ending of human transmission [5, 6]. However, in 2012, several cases of GW in dogs were reported in Chad along the Chari River [7] which prompted GWEP to include active surveillance of dogs and other animals in endemic areas. The number of reported infections in dogs increased significantly due to active surveillance and the provision of cash rewards for reporting suspected cases and infections [8]. Further population genetic studies confirmed that the emerging worms from dogs were *D. medinensis*, and that the same population of worms infect humans and dogs [9, 10]. A total of 886 animal infections were reported in 2023, the majority of them from dogs (The Carter Center News & Events 2024) [4]. Dogs in high-transmission areas along the Chari River are likely also infected through ingestion of frogs as paratenic hosts or fish as transport hosts carrying infected copepods [11–13].

In the absence of an effective therapy or vaccine, GW intervention is based on containment or tethering of the infected dogs and treating water resources with organophosphate larvicide temephos Abate^®^ to decrease copepod burden [7, 14]. However, considering the logistical challenges of applying these methods and their level of effectiveness, new tools are needed for the early detection of animal reservoirs to aid GWEP towards and beyond certification of countries as free of GW. Given the long prepatent period of 10-14 months for GW [1, 15], refined diagnostic methods are needed to detect infected animals for further containment or treatment if therapeutics become available in the future. Diagnostic tools will be essential to confirm the absence of GW transmission in animal reservoirs before further certification of remaining endemic countries. To find immunoreactive antigens, GW crude proteins were screened using plasma samples from infected humans and two proteins, thioredoxin-like protein 1 (TRX) and a domain of unknown function protein 148 (DUF148) were found [16]. These two antigens were further validated using a limited number of dog serum samples with recent GW infections from GW endemic areas in a multiplex bead assay [17]. While these two antigens showed promising results in detecting positive samples, they have not been assessed in detecting prepatent infections or evaluated for possible cross-reactivity with other parasitic nematodes.

In this study, we further validated these two GW antigens for accuracy in detecting GW infection using sera from experimentally infected dogs and ferret along with sera from Chadian dogs with a history of GW emergence. To determine possible cross-reactivity with other canine parasitic nematodes, we validated these antigens using sera from animal shelter dogs in Texas, United States, a non-endemic area for GW, using an indirect enzyme-linked immunosorbent assay (iELISA). We found that while DUF148 is more reactive and can detect prepatent infections, it cross-reacts with several sera from dogs from the GW non-endemic US. To mitigate cross-reactivity with DUF148, we produced shorter peptides and found that peptide 3 from the C-terminal site of the protein, is more immunogenic and showed the best performance for detecting GW infection. However, it did not improve the performance of whole antigen in the iELISA. Altogether, these results show the feasibility of DUF148 for detecting prepatent infections in the field.

## Methods

### Dog sera

#### GW endemic area

From September 2021 to May 2023, serum samples were collected during an intervention of the Guinea Worm Eradication Program in 56 villages along Chari River from three different regions (Moyen-Chari, Chari Baguirmi, and Mayo-Kebbi Est) of Chad, Africa. This study is part of a project to assess the impact of flubendazole treatment on the incidence of GW in dogs. All procedures were in accordance with the National Bioethics Committee of Chad (Protocol #005/PR/MESRI/SE/DGM/CNBT/SG/2022) and the University of Georgia’s Institutional Animal Care and Use Committee (A2019 04-005-Y4 A2). Initially, 1210 dogs were enrolled in the study in 2021 and more dogs were recruited in 2022, bringing the total number of dogs enrolled to 2495. Owner consent was obtained for all animals prior to enrollment and collection of samples. Blood was collected from dogs via venipuncture by trained personnel and then processed to separate serum. Serum samples were frozen, shipped to Texas A&M University, and kept at -80°C until processing. During this period, any GW emergence from enrolled dogs were recorded. A sample of the worm was sent to CDC for further confirmation of GW as per the established program guidelines.

#### Non-endemic GW area

Matching blood and serum samples were collected from 492 shelter dogs in Brazos and Harris counties, Texas, USA (IACUC 2022-0261 CA). These samples were originally collected for development of a probe-based qPCR assay for heartworm, *D. immitis* diagnosis [18]. Serum samples were screened for heartworm antigen using DiroCHEK^©^ (Zoetis Inc., Kalamazoo, MI, USA) which is a commercially available ELISA-based detection test [19]. Sera from dogs with either suspected or confirmed *O. lupi* infection were collected from New Mexico, USA, in another study to develop diagnostics for this parasite [20].

### Experimental animal infections

Two dogs and a ferret were experimentally infected with GW. All animal work and experiments were conducted in accordance with the regulations of the Animal Welfare Act, and the protocol was approved by the Texas A&M University Institutional Animal Care and Use Committee (AUP 2023-0273).

For dog infections, lab-reared copepods originally collected from Chad and identified as *Mesocyclops*, *Thermocyclops*, and *Eucyclops* through analysis of partial *cytochrome oxidase subunit 1* gene sequence [21, 22], were exposed to L1 GW larvae recovered from *D. medinensis* female worms removed from an experimentally infected ferret at the University of Georgia [23, 24]. The exposed copepods were maintained for 2 weeks to allow GW larvae to molt to the infective L3 stage. Two laboratory-raised female spayed beagles (Ridglan Farms, Inc., Mount Horeb, WI, USA) were each exposed *per os* to 40 copepods infected with GW L3 larvae. Dogs were group housed and provided food and water ad libitum. Blood samples were collected from the saphenous vein before inoculation and then weekly after GW-exposure. Whole blood and serum samples were frozen at −80°C until testing.

The ferret was infected with copepods exposed to L1 larvae from a gravid GW recovered from a naturally infected dog in Chad and transferred to Texas A&M University. The copepods were monitored for 18 days when they were confirmed to contain L3 stage GW larvae. Copepods were dissected under a microscope to release L3 larvae. A single female ferret was exposed intraperitoneally to 9 L3 larvae. Blood samples were collected before inoculation and then biweekly after inoculation.

### GW recombinant proteins production and peptides synthesis

Recombinant proteins were produced in *E. coli* by Biomatik (Biomatik, Canada). TRX and DUF148 are signal peptide-containing proteins and the recombinant proteins were expressed as mature protein without a signal peptide. Recombinant proteins were expressed as tag-free and purified using size exclusion chromatography. The lyophilized proteins were reconstituted in molecular grade water at a concentration of 0.5 mg/ml and kept in -80°C until use.

Three peptides were synthesized by Biomatik that covered full length of DUF148 with partial overlap between peptides (indicated in bold letters): Peptide 1: QFDDDIPPFLKGAPQSTIKEFETILQNGQSQTDQQLDA**NINAWIAKQTSA;** Peptide 2: **NINAWIAKQTSA**IQNAYRTFMAQIRTAQQQAEQARRTML**AKFSADARAAD;** Peptide 3: **AKFSADARAAD**AQLTKIAEDPRLTGEQKQAKIEATFKGSKLSNYNSRKCL. The peptides were reconstituted in DMSO at a concentration of 5 mg/ml and kept at -80°C until use.

### ELISA optimization

The microplates (Nunc Maxisorp, Thermo Fisher Scientific, Rockford, USA) were coated overnight at 4°C with 100 µL of 500 ng of GW proteins or 200 ng of DUF148 peptides diluted in 50 mM carbonate/bicarbonate buffer, pH 9.6. The wells were blocked with 5% skimmed milk in phosphate buffered saline with 0.05% Tween 20 (PBS-T), and the antigen-coated wells were filled with the diluted serum and incubated overnight at 4°C. The test sera (100 µL) were diluted 1:1000 with PBS-T containing 5% skimmed milk and while the secondary antibody (100 µL) was diluted in 1:10,000. Horseradish peroxidase (HRP) conjugated rabbit anti-dog IgG (Sigma, A6792) or goat anti-ferret IgG (Novus Biologicals, NB7224) was used as secondary antibody. Optical density (OD) was measured at 450 nm using Synergy H1 microplate reader (Biotek, Winooski, VT, USA). All the tests were done in duplicates.

### Statistical analysis

The ELISA sensitivity and specificity, and the optimal density cut-off, were calculated by plotting the receiver operating characteristic (ROC) curves using Prism. The area under the ROC curve (AUC) was estimated by non-parametric integration to measure diagnostic accuracy. To test for the statistical significance between mean OD values of GIN positive and negative sera, Mann-Whitney *U* test was performed. The values were considered significantly different if *P*<0.05.

## Results

### Optimization of GW antigen concentration for indirect ELISA (iELISA)

To find the best concentration of antigens for iELISA, we used a panel of serum samples from Chad dogs with history of GW emergence, shelter dogs from a non-endemic GW area, Texas (USA), and naïve, purpose-bred, laboratory-reared, beagle dogs. The plates were coated with 0.1 to 1000 ng of each GW recombinant protein (TRX and DUF148) or a cocktail of both proteins (1:1). We found that 500 ng of each antigen gave the best optical density (OD) difference between positive sera and negative sera (S1 Fig) and this concentration was used for coating the subsequent plates.

### iELISA of experimentally infected dogs and ferret with GW

We used the iELISA with each GW antigen or a cocktail of both antigens to screen the sera of two dogs (BEW and BDW) and a ferret experimentally infected with GW (Fig 1). The dogs were exposed via oral gavage with copepods infected with L3, and the ferret was exposed intraperitoneally to L3s that were removed from infected copepods. Sera from BEW showed increased reactivity at 9- and 11-weeks post-exposure to TRX and DUF148, respectively. However, the titer to both antigens gradually declined up to 14- month post-exposure when the necropsy was performed. No GW was retrieved from these dogs indicating that the infection did not establish. Ferret sera showed increased reactivity to TRX and DUF148 at 13- and 15-weeks post-exposure, respectively (Fig 1). The titer decreased gradually and had two more peaks at 21- and 49-weeks post-exposure, likely reflecting GW migration through tissues. The prepatent infection was confirmed by detection of subcutaneous worm through ultrasound at 9-months post- exposure (S2 Fig). Seven gravid female nematodes were retrieved from the ferret during necropsy at 12-months post-exposure (S1 Table).

**Fig 1.**
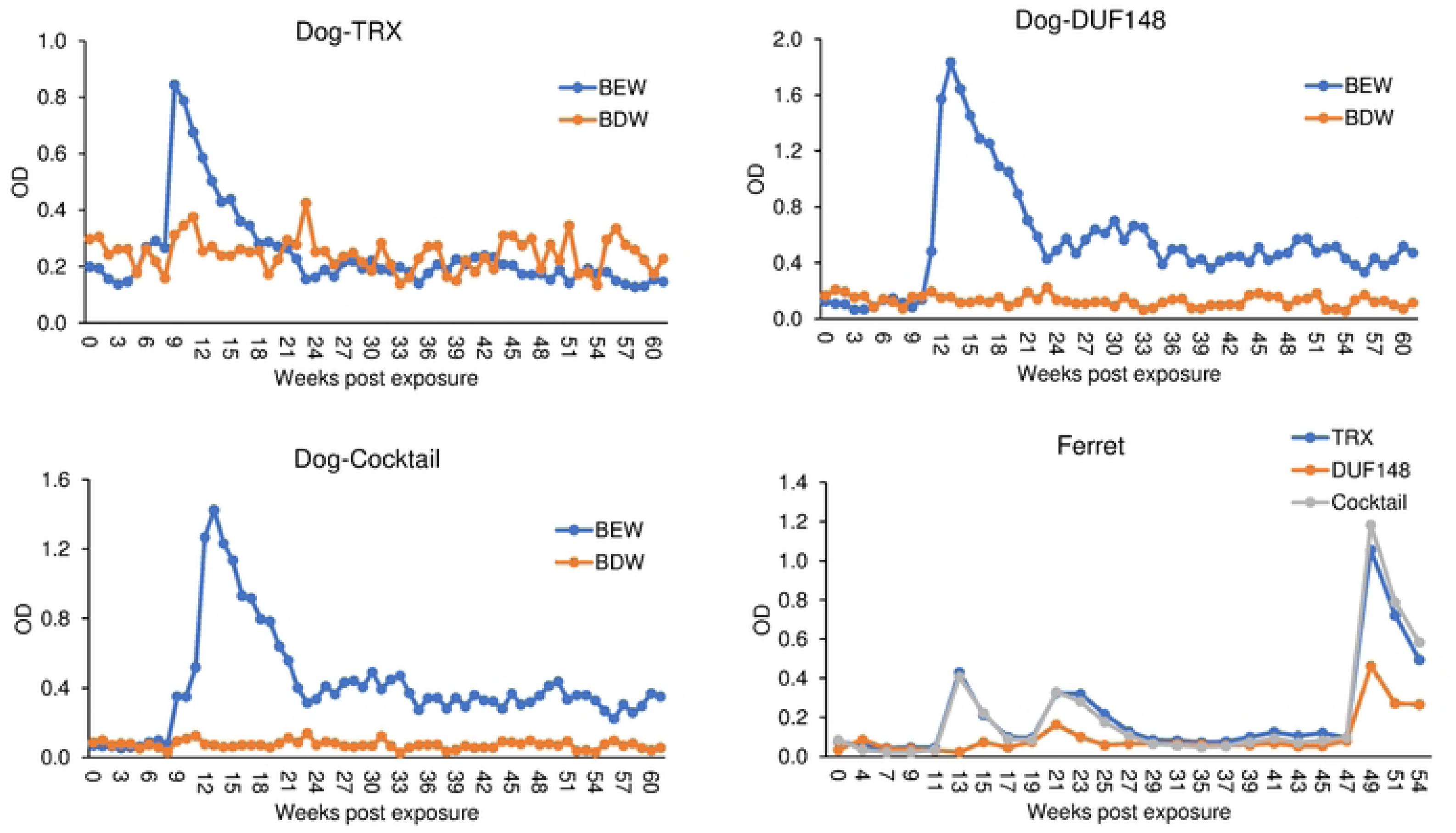
Time course reactivity of weekly (dogs) or biweekly (ferret) collected sera in TRX, DUF148 or cocktail iELISA. Dogs (BEW and BDW) were exposed to infected copepods with L3 stage and the ferret was exposed to L3 larvae at the timepoint 0.

### Evaluation of iELISA for detection of prepatent GW infection in Chad dogs

To validate the performance of DUF148 and TRX iELISA for detecting GW infection, archived sera from Chadian dogs with a history of GW emergence were selected. Considering 10-14 months prepatent period [1] and ∼4 month interval between ingestion of infected copepods and the appearance of antibodies in dogs based on our experimental GW infection, we selected 47 serum samples collected from dogs within 6 months prior to reported GW emergence. We used 492 US shelter dog sera as negative controls from a non-GW endemic area. The receiver operating characteristic (ROC) analyses were carried out to estimate the cut-off and performance indices, namely: area under the curve (AUC), sensitivity, specificity, positive predictive value (PPV), and negative predictive value (NPV) for DUF148 and TRX and the cocktail of both antigens (Fig 2, Table 1). DUF148 was more antigenic and showed a better performance than TRX with 76.6% sensitivity, 85.2% specificity, and an AUC of 0.87. The cocktail of both antigens showed a higher sensitivity (78.7%) but lower specificity (78.5%). However, 73 and 97 US dog serum samples cross-reacted with DUF148 and TRX, respectively. These dogs were screened for heartworm (*Dirofilaria immitis*) using different tests [20], and TRX and DUF148 cross-reacted with these sera regardless of heartworm positivity (Fig 3). TRX and DUF148 are signal peptide-containing proteins predicted to be secreted by GW and are conserved across the phylum Nematoda with considerable sequence similarity (S3 Fig). We screened DUF148 reactivity with sera from experimentally infected dogs with *Brugia pahangi*, *B. malayi*, and gastrointestinal nematodes (GIN; *Ancylostoma caninum*, *Toxocara canis*, *Toxascaris leonina*, and *Uncinaria stenocephala*). Cross- rection was observed with one *B. pahangi*-infected dog. We also assessed cross-reactivity with shelter dog sera for which paired fecal tests (double-centrifugal sugar flotation) were available, confirming GIN infection (S2 Table). We found that DUF148 reacted significantly more with the sera of dogs positive for GIN than sera of negative dogs (Fig 4). Additionally, we screened archived sera of US dogs that were naturally infected with the filarial nematode *Onchocerca lupi* for cross-reactivity with GW antigens. DUF148 cross-reacted with 4 samples while TRX cross-reacted with 14 samples (total 49 samples) (Fig 3). Overall, these results showed that GW antigens cross-reacted with sera of GIN- and *O. lupi-*positive dogs.

**Fig 2.**
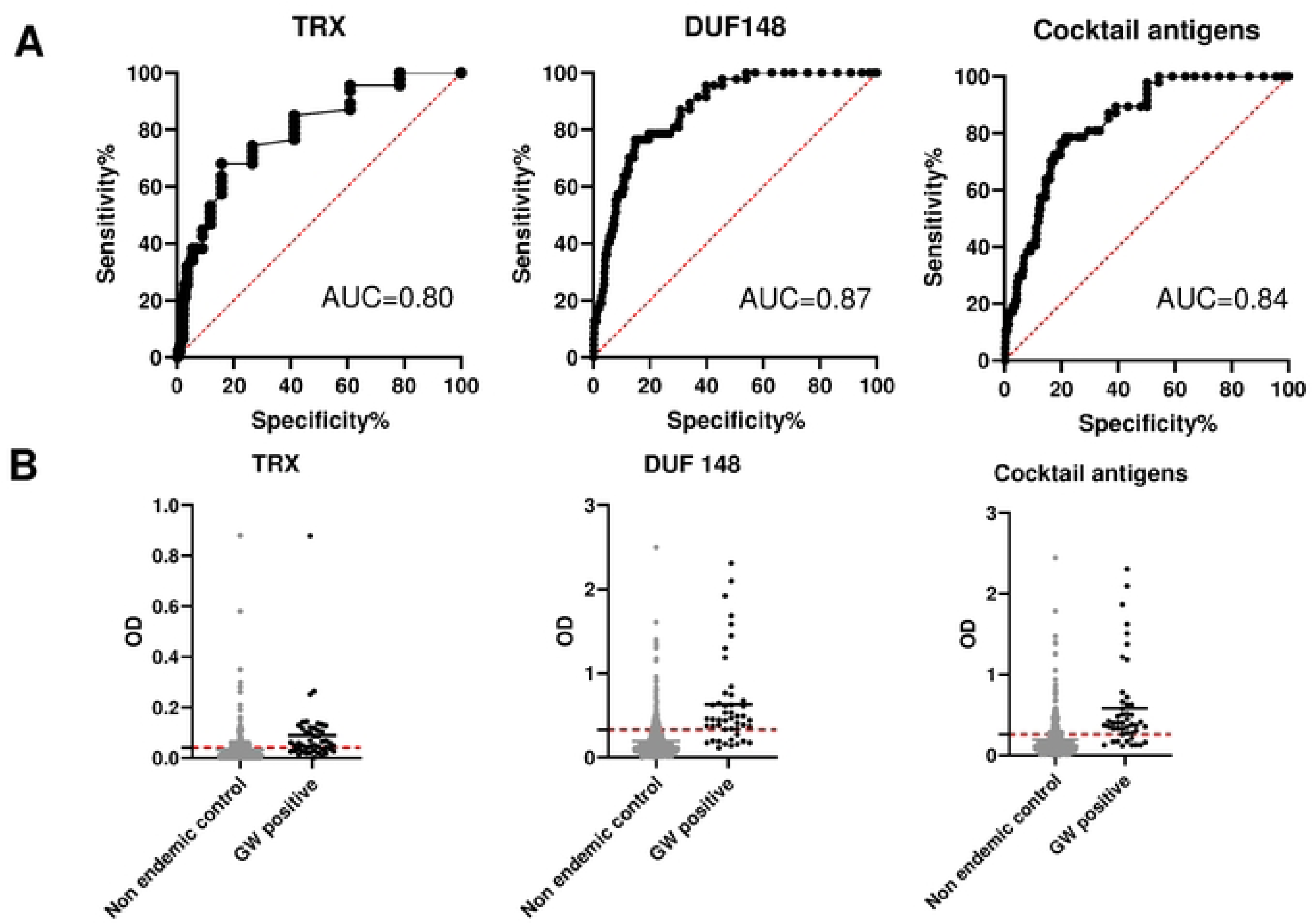
ROC analysis and reactivity of dog sera to GW antigens. **(A)** ROC analysis was carried out to define cut-off value, area under the curve (AUC), sensitivity, and specificity using 47 positive (Chad) and 492 negative (USA) dog sera. **(B)** Dashed lines represent the cut-off (determined by ROC curves) in the absorbance level.

**Fig 3.**
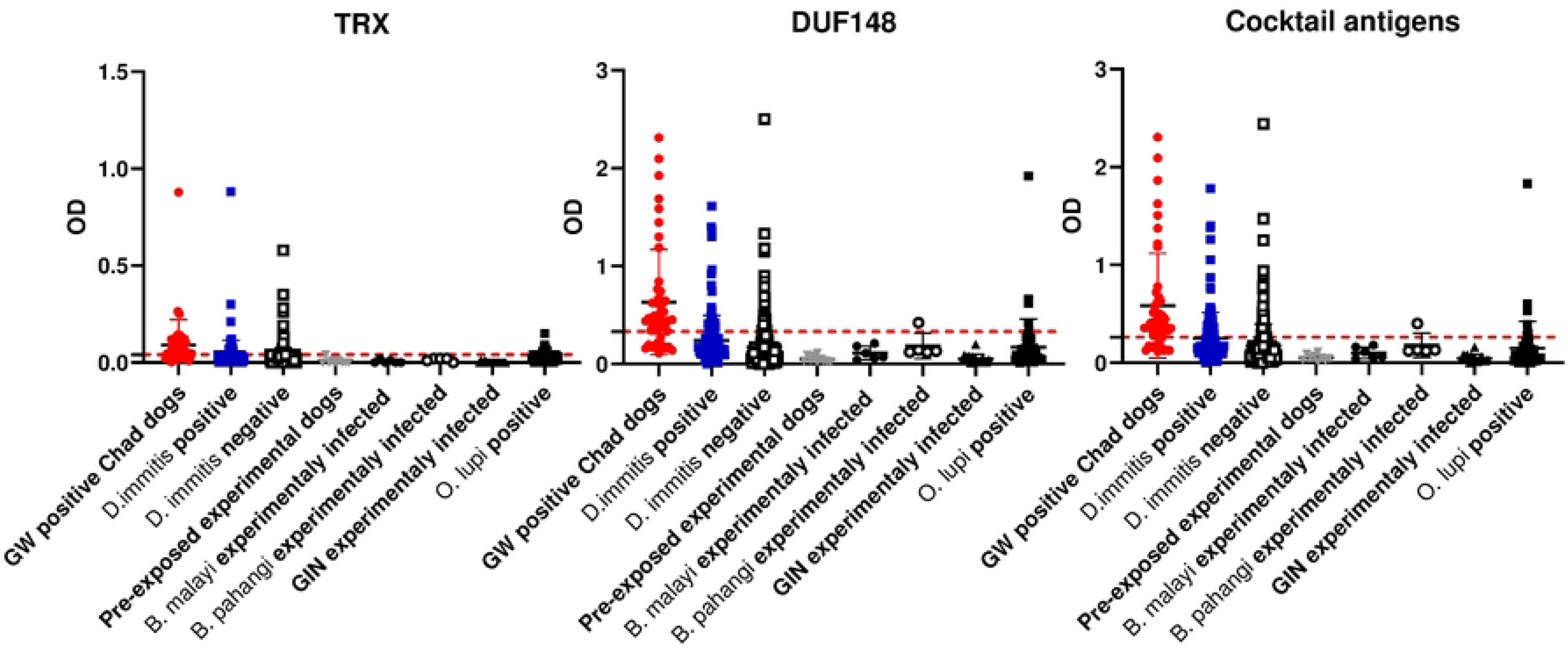
Optical density values for GW antigens among different groups of dogs. Mean OD values for each group are shown. Dashed lines represent the cut-off in the absorbance level determined by ROC curves. US shelter dogs are divided to *D. immitis* positive and negative groups. *D. immitis* and *O. lupi* dogs are natural infections.

**Fig 4.**
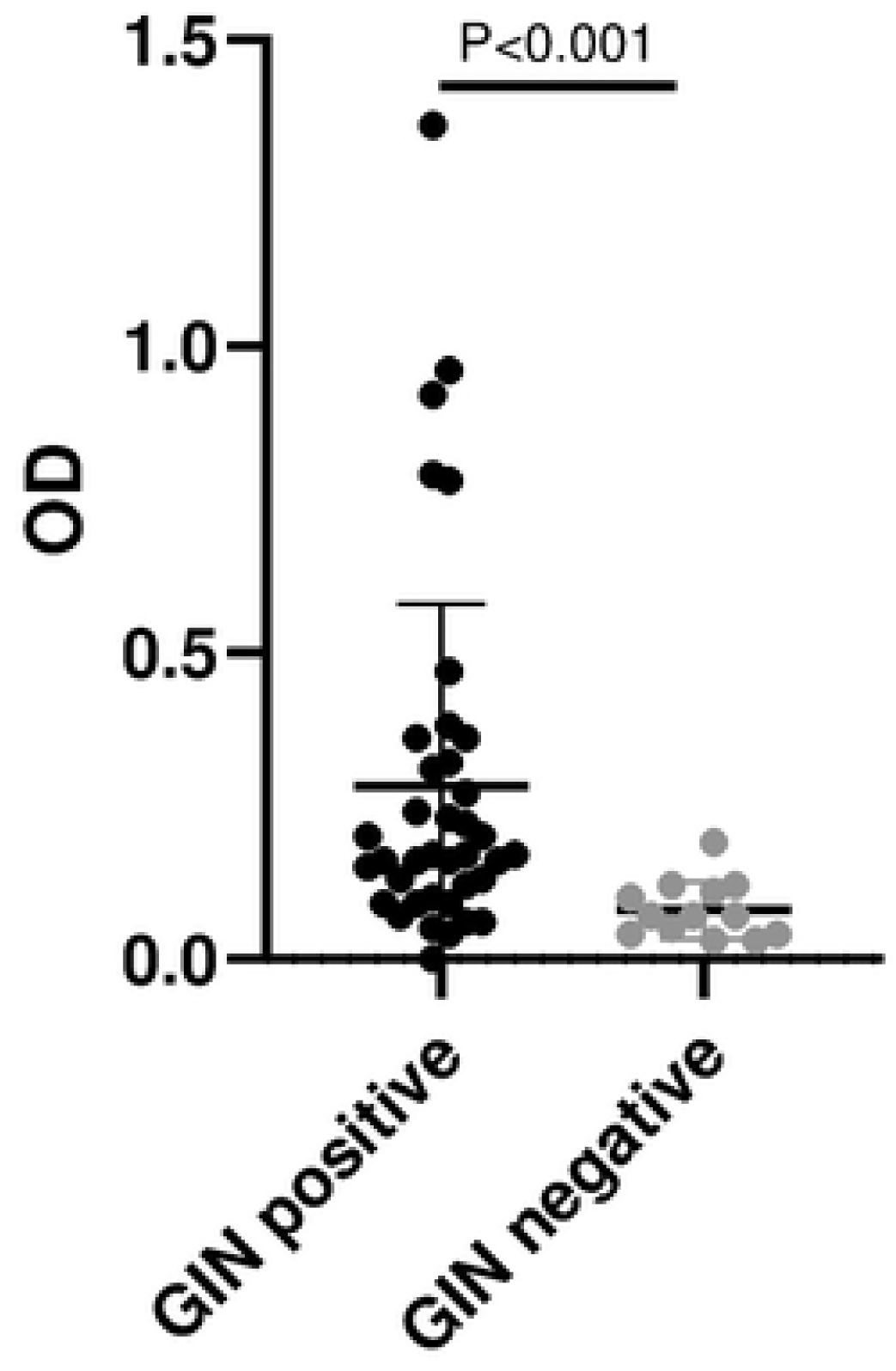
Reactivity of US shelter dog sera positive or negative for gastrointestinal nematodes (GIN) with DUF148. Thirty-eight fecal samples from shelter dogs were positive and 13 were negative for GIN eggs using double-centrifugal sugar flotation test. *P* value was determined by Mann-Whitney *U* test.

**Table 1.** Diagnostic performance of GW antigens and DUF148 peptides.

| Target Ag | AUC | CI | Cutoff OD | Sensitivity (%) | Specificity (%) | PPV (%) | NPV(%) |
| --- | --- | --- | --- | --- | --- | --- | --- |
| TRX | 0.80 | 95% | > 0.04025 | 68.1 | 84.4 | 68.1 | 80.6 |
| DUF148 | 0.87 | 95% | > 0.3313 | 76.6 | 85.2 | 76.6 | 86.5 |
| Cocktail Ag | 0.84 | 95% | > 0.2608 | 78.7 | 78.5 | 78.7 | 80.1 |
| Peptide 1 | 0.60 | 95% | < 0.09775 | 59.6 | 63.2 | ND | ND |
| Peptide 2 | 0.81 | 95% | < 0.06825 | 78.7 | 74.2 | ND | ND |
| Peptide 3 | 0.84 | 95% | > 0.1523 | 83 | 71.5 | 83 | 74.4 |
| Cocktail of P1&P3 | 0.84 | 95% | > 0.1003 | 85.1 | 69.3 | 85.1 | 69.8 |

AUC: area under the curve; CI: confidence interval; PPV: positive predictive value; NPV: negative predictive value.

### DUF148 peptide iELISA

To mitigate this cross-reactivity, we synthesized 3 short peptides from DUF148 that covered the whole mature DUF148 antigen. Each peptide partially overlapped with the other peptides. The concentration of each peptide to be used for iELISA was optimized (S4 Fig). Our initial screening confirmed that peptide 2 which constitutes the middle part of the protein was not immunoreactive to GW positive sera. Therefore, we proceeded with peptide 1 and 3 and a cocktail of peptides 1 and 3. Peptide 1 (the N-terminal of DUF148) was less reactive to GW positive sera than dog sera from the non-endemic US (Fig 5, Table 1). Peptide 3 which encompasses the C-terminal of DUF148 showed the best performance (Fig 5, Table 1) with sensitivity of 83%, specificity of 71.5%, and AUC of 0.84 in comparison to peptide 1 and did not cross-react with any sera from experimentally infected dogs with *Brugia* spp. or GIN (Fig 6). However, cross-reaction was observed with US shelter dogs and *O. lupi*-positive dogs. The cocktail of peptides 1 and 3 showed better sensitivity (85.1%) but the specificity declined (69.3%) in comparison to peptide 3 (Table 1). Overall, although peptide 3 showed a better sensitivity than DUF148 whole antigen but specificity and AUC were not improved.

**Fig 5.**
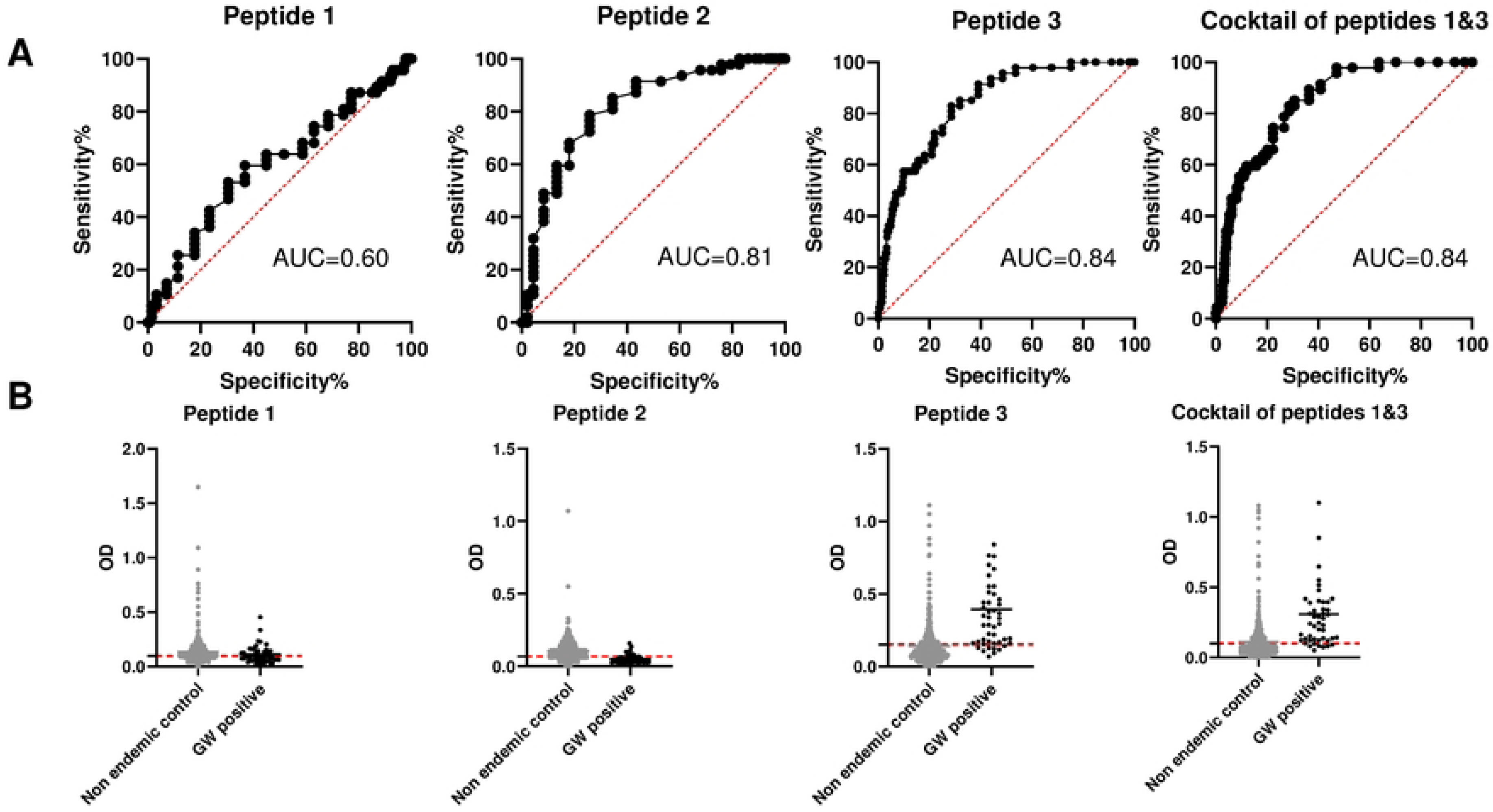
ROC analysis and reactivity of dog sera to DUF148 peptides. **(A)** ROC analysis was carried out using 47 positive (Chad) and 492 negative (USA) dog sera to define cut-off value, area under curve (AUC), sensitivity, and specificity. **(B)** Dashed lines represent the cut-off in the absorbance level.

**Fig 6.**
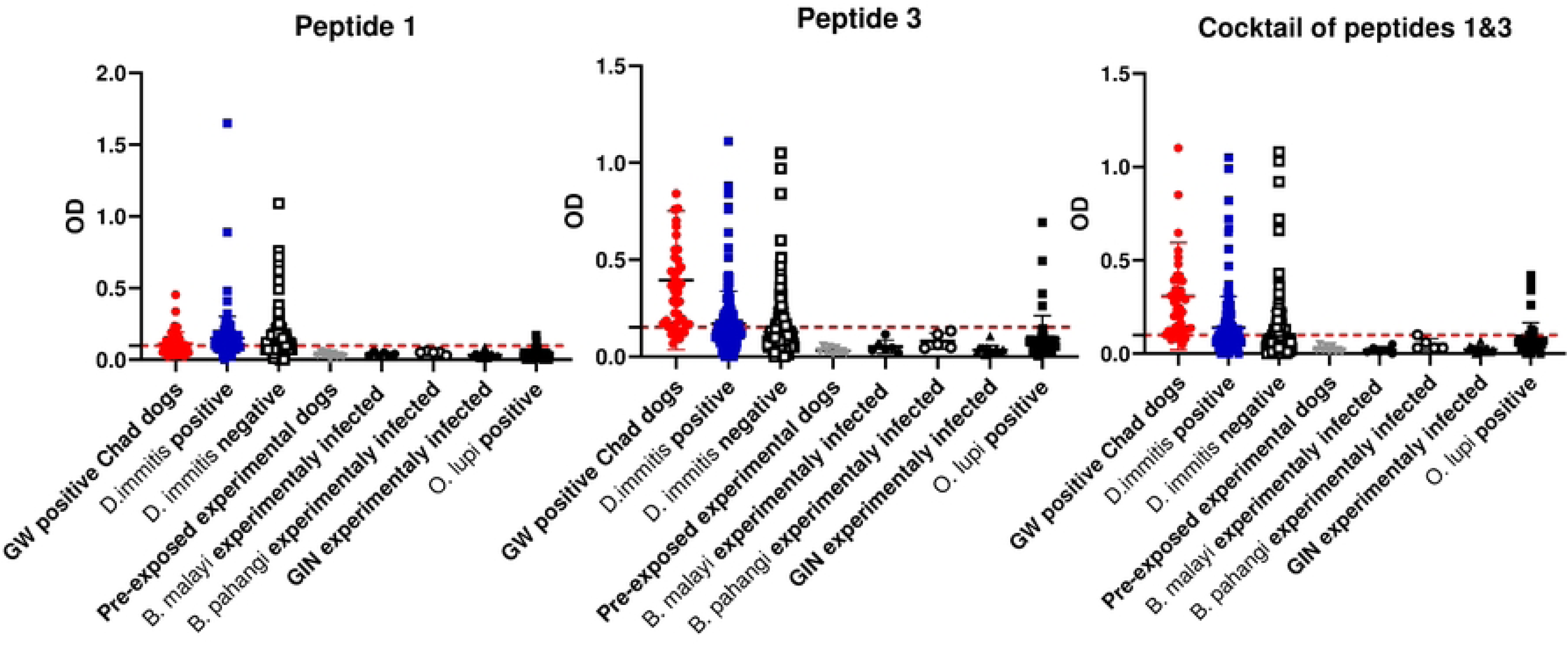
Optical density values for DUF148 peptides among different groups of dogs. Mean OD values for each group are shown. Dashed lines represent the cut-off in the absorbance level. US shelter dogs are divided to *D. immitis* positive and negative groups. *D. immitis* and *O. lupi* dogs are natural infections.

### iELISA of Chad dog sera pre- and post GW emergence

To test whether there is a correlation between antibody response to GW antigens and the days prior to GW emergence, we checked the reactivity to DUF148. We listed the dog sera within 6 months prior to GW emergence. Although we could observe a trend of increasing in the DUF148 titer to the day of worm emergence, there was no correlation between DUF148 titer and days of pre-GW emergence (Fig 7A). The antibodies to parasites antigens decay over time following the removal or clearance of the parasite. We checked the DUF148 titer in dog sera with single infection during the 6 months prior to GW emergence to two years after GW emergence. The dogs with multiple GW emergence events during the sampling period was defined as single infection if the worms emerged within a 1 month period. The highest titer was seen in the first 3 months post GW emergence and the titer waned over time up to 2 years post emergence (Fig 7B). However, anti-DUF148 titer was still detectable even after 2 years post GW emergence.

**Fig 7.**
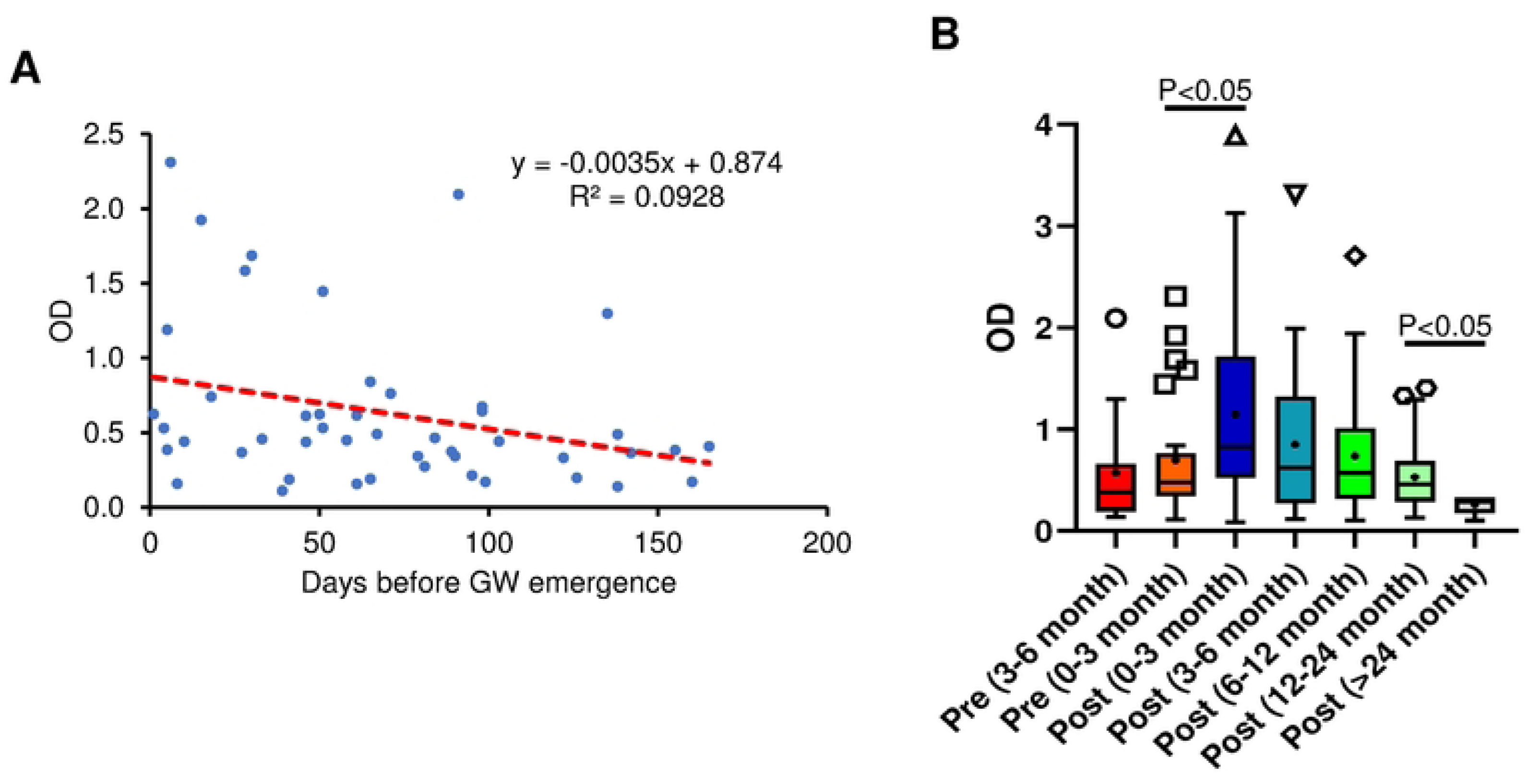
Correlation between antibody response to DUF148 and the time of GW emergence. **(A)** The DUF148 titer of dog sera (n=47) was plotted against days prior GW emergence. The dash line shows the linear correlation. GW emergence time is shown as 0 in the x axis. **(B)** Box plots denote median and interquartile ranges (whiskers denoting the median ± 1.5 X interquartile range) of different groups of dog sera (n=212) with GW emergence. *P* values are determined by Mann-Whitney *U* test.

## Discussion

Development of reliable diagnostic methods for detection of GW infection in reservoir animals is necessary to support GWEP efforts. Most animal infections in the remaining endemic countries are reported from domestic dogs. Detection of infected animals is necessary for further containment of animals to prevent spread of larvae into water sources or identifying animals for treatment when effective therapeutics become available in future. In this study we validated two GW antigens, TRX and DUF148, in iELISA format to be used for diagnosis of prepatent GW infection in dogs. These proteins were discovered as immunoreactive antigens with infected human plasma from endemic countries and were validated using a multiplex bead assay to detect the infection in dogs [16, 17]. In this prior study, both antigens were tagged with GST (glutathione S-transferase) which is derived from *Schistosoma* and used for purification of recombinant proteins [25]. Additionally, an in house-generated anti-dog IgG monoclonal antibody was used as commercial antibodies did not perform well in their multiplex bead assay [17]. TRX and DUF148 have signal peptides which indicates they are likely secreted by GW and could be promising targets for diagnosis. In our developed iELISA, we used mature proteins without tags to reduce the likelihood of cross-reactivity with other helminths, which would generate false-positive results. Additionally, a commercial anti-dog IgG was used as a secondary antibody. In experimental animals, specific antibodies against GW antigens were produced as early as 9 weeks post-exposure in dogs and 14 weeks in ferret. Anti-TRX antibodies appeared earlier than DUF148 in both animals indicating the different timing of these antigens release into host. Using Chad dog sera samples, we found that DUF148 is more reactive than TRX and was able to detect 76.6% (39 out of 47) of samples within 6 months prior to GW emergence. Target product profile of WHO for a serological test to detect prepatent GW infections in animals requires minimum 80% sensitivity and 90% specificity (https://www.who.int/publications/i/item/9789240090804).

Current sensitivity and specificity of DUF148 iELISA is 76.6% and 85.2%, respectively which is lower than the minimum required criteria. Our effort to use shorter peptides to mitigate cross-reactivity improved sensitivity of iELISA to 83% but could not improve specificity of the test.

DUF148 is a conserved antigen across Nematoda. To identify the possible reason for observed cross-reactivity with sera of dogs from the US, we screened several sera derived from dogs experimentally infected with different nematodes. We found that one *B. pahangi* infected serum sample reacted with DUF148. Dog infection with *Brugia* species has been reported from North America but not from Texas, USA [26]. Additionally, these shelter dogs were screened for common nematodes including heartworm and GIN [18, 19]. These sera reacted with DUF148 regardless of heartworm status. Additionally, we found that DUF148 reactivity is significantly higher in shelter dog sera positive for GIN than the sera of negative dogs. *Dracunculus insignis*, a common parasite of raccoons has been reported from dogs in Texas [27] and could be another contributing factor for cross-reactivity of US dog sera in our study with DUF148. Overall, these results suggest DUF148 likely cross-reacted with some GIN. GIN have a global distribution and it is likely that dogs in GW endemic regions, including Chad, are infected with these parasites. Additionally, a recent study showed the existence of *Brugia* sp. in Chadian dogs [28]. These GW antigens also cross-reacted with human sera of onchocerciasis patients (e.g., *Onchocerca volvulus*, *Wuchereria bancrofti,* and *Loa loa*) [16]. The results of our study and previous studies highlight the possibility of DUF148 cross-reaction with other nematodes.

No correlation was seen between antibody titer against DUF148 and days prior to GW emergence. Based on the antibody response to GW in ferret, it seems that the antibody titer fluctuates during the prepatent period and likely reflects GW migration through subcutaneous tissue. The titer significantly increased following GW emergence in Chadian dogs likely due to the local inflammation and the blister formation at the site of emergence. Although the antibody response waned gradually overtime, the anti-DUF148 antibody was detectable even at two years post emergence. Several factors may contribute to antibody response including the duration post infection, intensity of infection, and immune status of the host. Additionally, dogs in endemic GW regions are constantly at risk of infection, which could be the reason behind the prolonged antibody response. Our findings suggest that DUF148 is not a suitable antigen to track the antibody response following GW clearance.

In conclusion, DUF148 is a promising target for serological test to detect prepatent infection. In the absence of any diagnostic methods for animals in endemic countries, DUF148 iELISA could be a starting point for detection and management of infected dogs. TRX and DUF148 are valuable targets to track experimentally infected animals under laboratory conditions. However, finding more specific serological markers is needed for use in endemic regions under field conditions. Novel comprehensive methods to find serological markers are based on the proteome of the target organism. PepSeq technology, which is based on building a peptide library from the organism proteome [29], allows comprehensive screening of immunogenic linear peptides of GW. Following discovery of immunogenic antigens, the magnitude and longevity of IgG response should be checked to find antigens that can detect recent exposure. Serological tests would be useful to track host antibodies and whether GW is circulating in the animal population in endemic areas of the world.

## Supporting information

**S1 Fig. Optimization of GW antigen concentration for iELISA.** GW positive or negative (US shelter dogs and pre-immune sera from experimental dog, BEW) dog sera reactivity with different antigen concentrations (0.1-1000 ng) are plotted.

**S2 Fig. Experimental GW infection of ferret.** Subcutaneous worm was observed on the right side of ferret and GW was confirmed using ultrasound imaging at 9 months post exposure.

**S3 Fig. Multiple amino acid sequence alignment of GW DUF148 with its orthologs of other nematodes.** Asterisks show the conserved amino acids.

**S4 Fig. Optimization of DUF148 peptides concentration for iELISA.** GW positive or negative (US shelter dogs and pre-immune sera from experimental dog, BEW) dog sera reactivity with different antigen concentrations (0.1-1000 ng) are plotted.

## Acknowledgement

We would like to thank The Carter Center and the Programme National d’Éradication du Ver de Guinée, Ministry of Health, and the Afrique One ASPIRE, Institut de Recherche en Élevage pour le Développement, for technical and logistic support for field sample collections. We also thank various collaborators for providing serum samples from experimentally infected helminths, namely TRS, East Tennessee Clinical Research, Inc., the University of Georgia, and the NIH/NIAID Filariasis Research Reagent Resource Center (FR3) (www.filariasiscenter.org). We are grateful to The Carter Center staff for critically revising of our manuscript.

## Authors contribution

**Conceptualization:** Hassan Hakimi, Guilherme G. Verocai

**Data curation:** Hassan Hakimi, Guilherme G. Verocai

**Formal analysis:** Hassan Hakimi, Pabasara Weerarathne, Guilherme G. Verocai

**Funding acquisition:** Meriam N. Saleh, Lucienne Tritten, Guilherme G. Verocai

**Investigation:** Hassan Hakimi

**Methodology:** Hassan Hakimi, Pabasara Weerarathne, Richard Ngandolo, Philip Ouakou Tchindebet, Sidouin K. Metinou, Guilherme G. Verocai

**Project administration:** Hassan Hakimi, Guilherme G. Verocai

**Resources:** Meriam N. Saleh, Raquel R. Rech, Richard Ngandolo, Philip Ouakou Tchindebet, Sidouin K. Metinou, Lucienne Tritten, Guilherme G. Verocai **Supervision:** Hassan Hakimi, Guilherme G. Verocai

**Validation:** Hassan Hakimi, Guilherme G. Verocai

**Visualization:** Hassan Hakimi

**Writing – original draft:** Hassan Hakimi

**Writing – review & editing:** Hassan Hakimi, Pabasara Weerarathne, Meriam N. Saleh, Raquel R. Rech, Richard Ngandolo, Philip Ouakou Tchindebet, Sidouin K. Metinou, Lucienne Tritten, Guilherme G. Verocai

